# APEX-based proximity labeling in *Plasmodium* identifies a membrane protein with dual functions during mosquito infection

**DOI:** 10.1101/2020.09.29.318857

**Authors:** Jessica Kehrer, Dominik Ricken, Leanne Strauss, Emma Pietsch, Julia M. Heinze, Friedrich Frischknecht

## Abstract

Transmission of the malaria parasite *Plasmodium* to mosquitoes necessitates gamete egress from red blood cells to allow zygote formation and ookinete motility to enable penetration of the midgut epithelium. Both processes are dependent on the secretion of proteins from distinct sets of specialized vesicles. Inhibiting some of these proteins has shown potential for blocking parasite transmission to the mosquito. To identify new transmission blocking vaccine candidates, we defined the microneme content from ookinetes of the rodent model organism *Plasmodium berghei* using APEX2-mediated rapid proximity-dependent biotinylation. Besides known proteins of ookinete micronemes, this identified over 50 novel candidates and sharpened the list of a previous survey based on subcellular fractionation. Functional analysis of a first candidate uncovered a dual role for this membrane protein in male gametogenesis and ookinete midgut traversal. Mutation of a putative trafficking motif in the C-terminus led to its mis-localization in ookinetes and affected ookinete to oocyst transition but not gamete formation. This suggests the existence of distinct functional and transport requirements for Plasmodium proteins in different parasite stages.

**Significance:** The genome of the malaria parasite *Plasmodium* contains over 5500 genes, of which over 30% have no assigned function. Transmission of *Plasmodium spp*. to the mosquito contains several essential steps that can be inhibited by antibodies or chemical compounds. Yet few proteins involved in these processes are characterized, thus limiting our capacity to generate transmission interfering tools. Here, we establish a method to rapidly identify proteins in a specific compartment within the parasite that is essential for establishment of an infection within the mosquito, and identify over 50 novel candidate proteins. Functional analysis of the top candidate identifies a protein with two independent essential functions in subsequent steps along the *Plasmodium* life cycle within the mosquito.

**Highlights:**

- first use of APEX based proximity ligation in Apicomplexa
- identification of >50 putative ookinete surface proteins
- novel membrane protein essential for microgamete egress and ookinete migration
- putative trafficking motif essential in ookinetes but not gametes

## Introduction

The causative agents of malaria, unicellular *Plasmodium* spp., survive on a complex life cycle between a vertebrate and a mosquito vector. Interruption of the life cycle is a common goal of intervention studies (Coelho et al., 2019; Goh et al., 2019; Yahiya et al., 2019). To this end proteins on the parasite surface can in principle be targeted by antibodies or drugs; yet many of these proteins have neither not been identified or nor been characterized in depth. Life cycle progression of the parasite and surface location of proteins relies on the secretion of specific vesicles, which differ in content and size from one parasite stage to another. Distinct vesicles are important for intracellular stages to escape from their host cells or the oocyst and for extracellular stages to migrate, cross tissue barriers and enter into host cells. When an *Anopheles* mosquito feeds on an infected individual, it takes up infectious sexual parasite stages, so-called gametocytes, during the blood meal. Within the mosquito gut they rapidly differentiate into female macrogametes and male microgametes that both rely on the release of egress vesicles to escape from their surrounding infected red blood cells (iRBCs). Successful and efficient egress is essential for male gametes to fertilize females and form zygotes which develop into ookinetes. (Bennink et al., 2016; Flieger et al., 2018). Ookinetes, like red cell infecting merozoites and liver cell infecting sporozoites, depend on the discharge of micronemes for motility, which enables ookinetes to pass through the midgut epithelium to form an oocyst (Angrisano et al., 2012). While gametocyte egress vesicles appear randomly distributed in the non-polarized cell, micronemes are located at the apical end of the highly polarized migrating parasite forms (Dubremetz et al., 1998). Micronemes contain soluble and membrane bound proteins including adhesins, which are needed for attachment of the parasite to host tissue and cells, a prerequisite for motility and invasion. Known micronemal proteins can function in just one (e.g. CTRP, TRAP) or several parasite stages (e.g. GEST, PAT), while others are expressed at multiple stages but appear to only function during one (e.g. MTRAP) (Bargieri et al., 2016; Kehrer, Frischknecht, et al., 2016; Klug et al., 2018; Talman et al., 2011).

Our limited functional understanding of these proteins suggests that many more proteins are likely to play essential roles in parasite egress, adhesion, motility and invasion (Lal et al., 2009; Lindner et al., 2013). Yet, classic proteomic approaches often suffer from unspecific contaminations. For example density gradient centrifugation to enrich ookinete-derived vesicles containing CTRP, resulted in a list of 330 putative micronemal proteins (Lal et al., 2009). This list included several ribosomal and cytoskeletal proteins, which are most likely cytoplasmic contaminants of the enrichment procedure. To more specifically investigate vesicular content, we recently used the promiscuous biotin ligase BirA* (Roux et al., 2012) coupled to the *Plasmodium berghei* gametocyte specific protein MDV/PEG3 (Kehrer, Frischknecht, et al., 2016). This allowed enzyme-based proximity labeling of proteins in egress vesicles of living parasites with biotin (BioID). The labeled proteins could be purified via streptavidin coated beads and identified with mass spectrometry. From the list of the obtained proteins we localized several ones via fluorescent protein tagging to the secretory vesicles of gametocytes and revealed the involvement of MTRAP in gametocyte egress (Kehrer, Frischknecht, et al., 2016). BirA*-mediated BioID has since been used to identify new components of the parasitophorous vacuolar membrane, the apicoplast, rhoptries and Kelch13 interaction partners of blood stages (Geiger et al., 2020; Birnbaum et al., 2020; Boucher et al., 2018; Khosh-Naucke et al., 2018; Schnider et al., 2018) (Fig. 1A). However, one drawback of BirA*-mediated proximity labeling is the necessary long reaction time of up to 24h (Figure 1B). This did not allow us to identify the vesicular content of ookinete micronemes and motivated us to use the much faster labeling APEX2, a peroxidase isolated from soybean (Figure 1B). While BirA* catalyzes biotinylation of free amino groups, APEX2-mediated biotinylation occurs within just a minute and prefers electron rich amino acids such as tyrosine (Lam et al., 2014; Samavarchi-Tehrani et al., 2020).

**Fig. 1.**
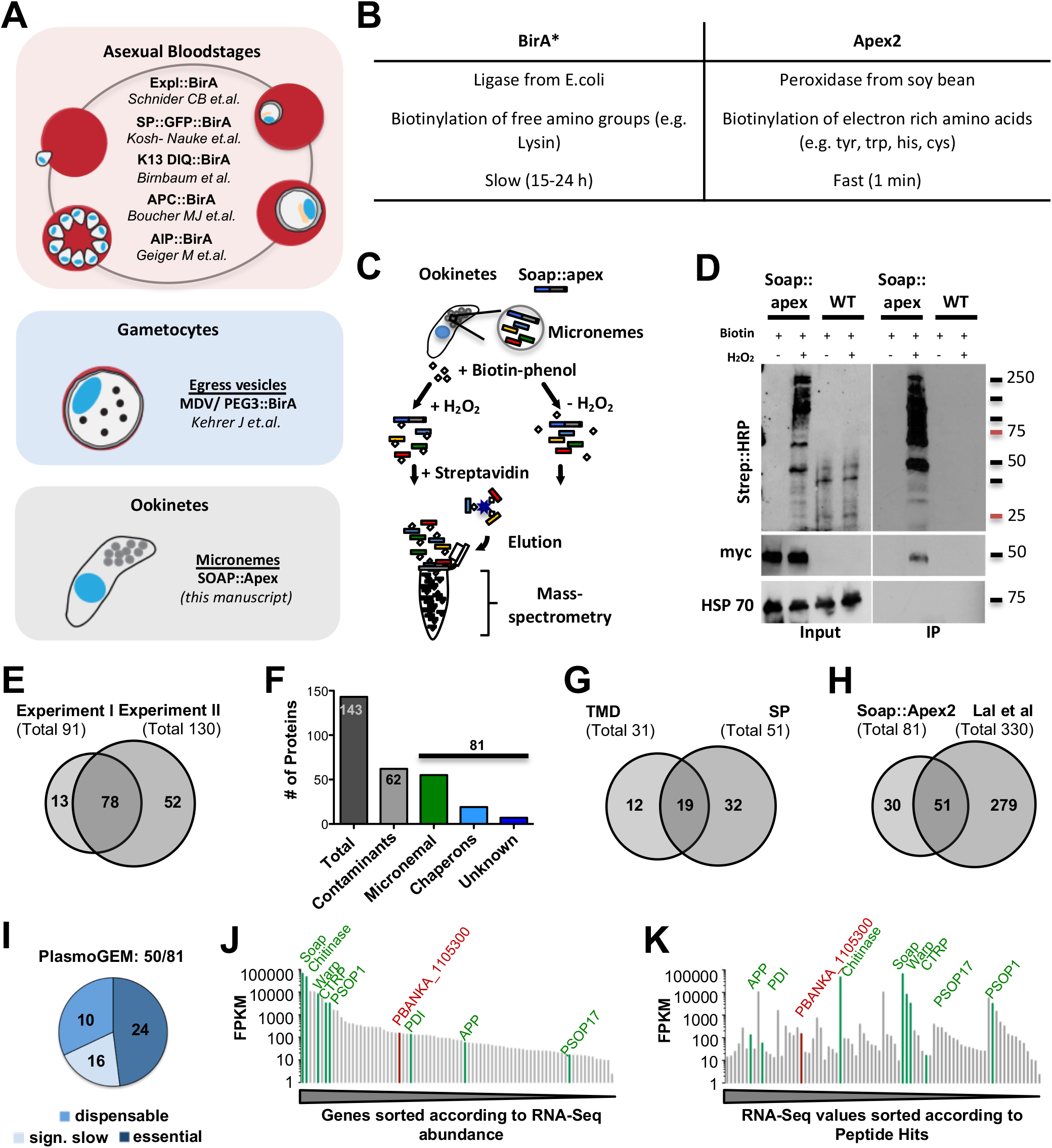
Proximity labeling in *Plasmodium* and in ookinetes. **(*A*)** Overview of BioID approaches performed in *Plasmodium* parasites. **(*B*)** General comparison of the BirA* and APEX2 based proximity labeling methods. **(*C*)** Schematic of APEX2 labeling for ookinete micronemes and sample processing for protein enrichment and mass-spectromerty analysis. **(*D*)** Western blots showing samples of wild type and *soap::APEX2* parasites before (input) and after (IP) enrichment of biotinylated proteins, with and without induction of labeling (H_2_O_2_). **(*E*)** Two experiments yielded 91 and 130 proteins with at least 2 peptide hits. 78 proteins were found in both experiments. **(*F*)** Of 143 identified unique proteins 81 are likely micronemal proteins while 62 were classified as likely contaminants (see table S1). **(*G*)** 31 identified micronemal candidate proteins contained at least one trans-membrane domain (TMD) and 51 a signal peptide (SP), 19 contained both. **(*H*)** 51 of the proteins identified in our BioID screen were also found by a subcellular fractionation approach (Lal et al., 2009). **(*I*)** 50 of the 81 micronemal candidate proteins were already investigated by the PlasmoGEM screen of which 24 were classified as essential, 16 conferred slow growth upon deletion and 10 were dispensable. **(*J*)** The top candidate, PbANKA-1105300 (red) by peptide hits that contained a SP and at least one TMD and was not characterized by the PlasmoGEM screen. **(*J-K*)** RNA-seq abundance (y-axis) of the identified micronemal candidate proteins (Otto et al., 2014) with known proteins (green) and the top candidate, PBANKA_1105300 (red) marked sorted (x-axis) according to RNA-seq data **(L)** and by peptide abundance from our BioID screen **(K)**.

Fusing the micronemal ookinete protein SOAP with APEX2 allowed the robust identification of known micronemal proteins of *Plasmodium* ookinetes as well as a list of unknown proteins.

We found 19 proteins that contained both a signal peptide and at least one transmembrane domain. We investigated the top hit in this category, a protein conserved within *Plasmodium* spp. featuring four transmembrane domains and a long divergent extracellular coiled-coil domain by fluorescent tagging, gene deletion and complementation with the *P. falciparum* orthologue. This showed that the protein, termed akratin (from the Greek for powerless/impotent), is expressed in nearly all stages along the parasite life cycle and appears essential for microgametogenesis as well as for efficient ookinete migration. Mutation of a putative micronemal targeting motif in the C-terminus of the protein impacted ookinete migration through the midgut epithelial but not gamete egress, suggesting distinct trafficking requirements for targeting proteins to different types of vesicles.

## Results

To identify micronemal proteins of ookinetes we used the microneme-resident protein SOAP (Secreted Ookinete Adhesive Protein, PBANKA_1113400) as bait for proximity labeling. As ookinetes are fully formed within 21 h, but proximity based biotinylation with BirA* may require more than 24 h of incubation, we turned to the faster-acting ascorbate peroxidase APEX2 (Lam et al., 2014). SOAP is a small protein of 166 amino acids with a molecular weight of around 21 kDa and important for ookinete to oocyst transition (Dessens et al., 2003). In analogy to our previous approach (Kehrer, Frischknecht, et al., 2016) we generated a parasite line expressing SOAP C-terminally fused to APEX2 and a 3x myc tag (Figure S1). Fully developed and purified SOAP::APEX2 and wild type control ookinetes were incubated with biotin-phenol for 30 min at room temperature. The cells were then split into two separate populations and biotinylation was initiated in one vial for 1 min using H_2_O_2_ while the second vial served as control (Figure 1C). Samples before and after enrichment of biotinylated proteins with streptavidin coated beads were tested for successful labeling of proteins by western blot (Figure 1D) and used for proteomic analyses.

In two independent experiments we identified 143 potential micronemal proteins of which 78 appeared in both experiments (Figure 1E, Table S1). Of these 143 proteins, 62 proteins were likely not micronemal such as the cytoskeletal proteins actin and tubulin, ribosomal proteins and those resident in the Golgi. Hence, we classified them as likely contaminants (Figure 1F, Table S1). Among the remaining 81 putative microneme-resident proteins for which at least two peptides were detected, we found the well-known micronemal ookinete proteins CTRP, WARP, SOAP, chitinase, PSOP17 and PSOP1 (Dessens et al., 2001; Dessens et al., 1999; Dessens et al. 2003; Ecker et al., 2008; Yuda et al., 2001) (Table S1) as well as aminopeptidase (APP) and protein disulfide isomerase (PDI), which have been localized to vesicles reminiscent of micronemes but not functionally studied in ookinetes (Ecker et al., 2008; Ghosh et al., 2018; Lal et al., 2009). Furthermore, we identified 19 chaperones and seven conserved *Plasmodium* proteins with unknown function, which we listed as likely micronemal candidates (Figure 1F, Table S1). The top 20 candidate proteins with the most hits (over 20 peptides) contained 19 putative micronemal proteins including four chaperones and only one contaminant (alpha tubulin 2) with the three most abundant proteins (99 to 135 hits) being members of the LCCL domain containing protein family (CCp/ Lap) (Saeed et al., 2010). The 54 candidate proteins with 5 to 20 peptide hits already featured 52% likely false positives and four chaperones while in the 69 proteins with only two to four peptide hits we found a similar number of contaminants, 48%, with the number of chaperones increasing to 11 (Table S1). Yet, even this fraction contained known or likely micronemal proteins such as PSOP1 (Ecker et al., 2008) and the ookinete surface protein p25 (Saxena et al., 2007; Tomas et al., 2001) (Table S1).

51 of the down-selected 81 proteins contained an N-terminal signal peptide (SP) and 31 showed at least one transmembrane domain (TMD) with 19 proteins featuring both (Fig. 1G). Only 18 proteins did not show any detectable domain as annotated on PlasmoDB (Table S1). Comparing the 81 proteins with previous data from a cell fractionation study reporting 330 potential micronemal proteins in ookinetes (Lal et al., 2009) showed 51 common proteins but also 30 novel candidates (Figure 1H, Table S1). To aid candidate selection we next interrogated the PlasmoGEM database ((Schwach et al., 2015) plasmogem.org) for genes encoding proteins in our list that were targeted by the batch gene-deletion approach (Bushell et al., 2017). 24 of the 50 genes targeted by PlasmoGEM were reported to be essential for blood stage development, while parasites lacking 16 further proteins were growing significantly slower and 10 were reported as dispensable genes (Figure 1I, Table S1).

We finally ranked our identified 81 candidates to ookinete RNA Seq data (Otto et al., 2014) sorted by the number of peptide hits per protein (without taking protein length into account) or total RNA Seq abundance (Figure 1 J-K). While the known proteins CTRP, SOAP, WARP, chitinase and PSOP1 were the most abundant when sorted according to RNA Seq data, it was surprising that these proteins were not appearing among the top hits when sorted according to the number of detected peptide hits.

To investigate a first candidate protein from our list by functional analysis, we selected the protein with the highest numbers of peptide hits, containing both a SP and at least one TMD. This protein, PbANKA_1105300, contains 455 amino acids and is predicted to feature a signal peptide, four TMDs (as identified through TMHMM and plasmodb.org) and two coiled-coil domains (http://smart.embl-heidelberg.de), suggesting a role in protein-protein interactions (Figure 2A, Figure S1). PbANKA_1105300 was not present in our previous BioID screen from gametocytes and was not probed by PlasmoGEM or present in the study of Lal et al. 2009, but featured in the total sporozoite proteome (Lindner et al., 2013). Using BlastP (https://blast.ncbi.nlm.nih.gov/Blast.cgi) PbANKA_1105300 was found to be conserved among *Plasmodium* spp. It shares 82 % identity with its orthologue in *P. yoelii*, but only 35 % and 32 % with its orthologues in *P. falciparum* and *P. vivax*, respectively. In contrast to *P. berghei* and *P. falciparum*, the orthologues in *P. yoelii* and *P. vivax* are predicted to contain only two TMDs. Furthermore, the *P. falciparum* protein contains about two times more amino acids than the *P. berghei* protein. No orthologue was found outside *Plasmodium* spp. (Figure 2B, Figure S3). Transcriptome data of PbANKA_1105300 showed a peak abundance in fully developed ookinetes in comparison with lower expression in gametocytes, asexual stages and sporozoites (Figure 2C)(Otto et al., 2014). Comparing data of male and female gametocytes, expression was found to be higher in males (Lindner et al., 2013; Otto et al., 2014; Yeoh et al., 2017) (Figure 2D).

**Fig. 2.**
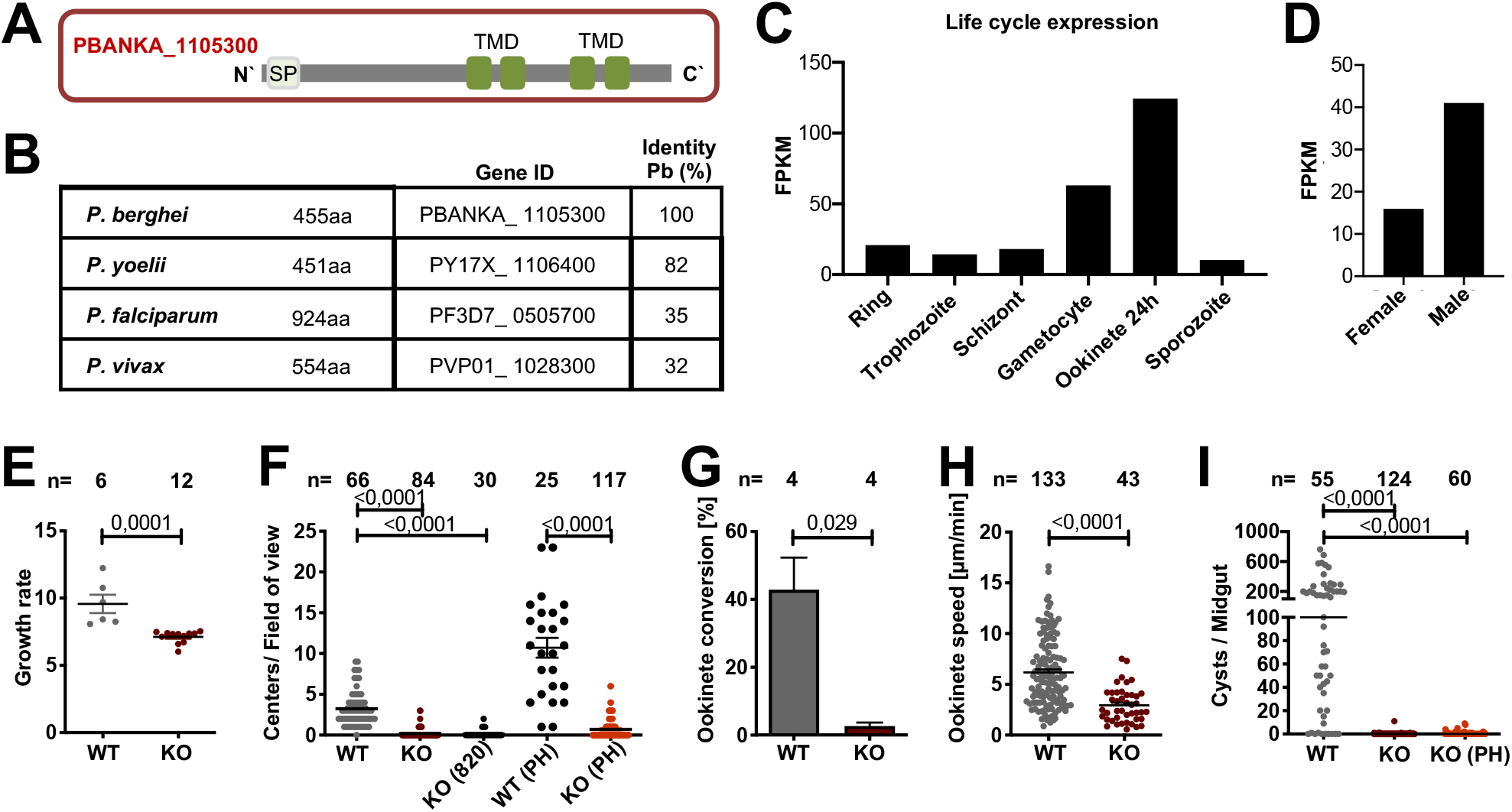
Akratin is expressed across the life cycle and essential for mosquito infection. **(*A*)** In *P. berghei* the protein contains a signal peptide (SP) and 4 transmembrane domains (TMD). **(*B*)** Akratin (PbANKA_1105300) is conserved across *Plasmodium spp. **(C-D)** Akratin* expression (RNA-seq values) across the life cycle **(*C*)** and in male and female gametocytes **(*D*);** (Lindner et al., 2013; Otto et al., 2014) **(*E*)** Blood stage growth rate of *akratin(-)* (KO) parasites in comparison to wild type. Data points represent parasites growing in individual mice. **(*F*)** Exflagellation of microgametes in the presence and absence of phenylhydrazin (PH) is nearly absent in KO parasites. **(*G*)** *Akratin(-)* parasites show severely reduced ookinete conversion. **(*H*)** Speed of gliding *akratin(-)* and wild type (WT) ookinetes. **(*I*)** Infected *Anopheles stephensi* mosquitoes barely show any midgut infections even in the presence of phenylhydrazin (PH). P-values calculated by Mann Whitney test (E, G, H) and Kruskal Wallis Test with Dunns multiple comparison (F, I). Horizontal bars indicate mean ± SEM.

### Parasites lacking *PbANKA_1105300* (akratin) fail to transmit to mosquitoes

To determine a possible function of PbANKA_1105300, we generated a *P. berghei* parasite line lacking the gene by replacing its open reading frame (ORF) with a selection cassette suited for fluorescent activated cell sorting (FACS) (Figure S3). Characterization of the resulting polyclonal parasites showed a significant reduction in blood stage growth from around 10 as observed for wild type parasites to seven for parasites lacking PbANKA_1105300 (Figure 2E). More strikingly, parasites lacking PbANKA_1105300 showed a strong reduction in male exflagellation, which could not be restored in phenylhydrazin (PH) treated mice. (Figure 2F). We hence named the protein akratin (Greek: akrates, for powerless/impotent). *Akratin(-)* parasites consequently showed reduced gametocyte to ookinete conversion rates (Figure 2G) and a slight reduction of ookinete motility (Figure 2H), although this might be due to the generally low numbers of ookinetes obtained in the cultures. Importantly, feeding of mosquitoes on infected mice failed repetitively to yield oocysts, hence revealing a complete block of the transgenic parasites in transmission to the insects (Figure 2I).

### Complementation with the *P. falciparum* orthologue rescues *akratin(-)* defects

To exclude a phenotype caused by off-target effects we next generated two parasite lines where we complemented the *akratin(-)* line with a GFP-tagged version of either *P. berghei* or *P. falciparum* (PF3D7_0505700) akratin, before analyzing the observed gene deletion phenotype in more detail. After recycling the selection cassette of the KO using negative selection (Figure S3) we inserted the respective gene into the selection marker-free parasites (Figure S4). Both parasite lines containing either the original *P. berghei* gene or the *P. falciparum* orthologue C-terminally tagged with GFP rescued the defect in gamete egress and the transmission block of the gene deletion mutant. Indeed, the observed infection rates of mosquitoes as well as oocysts, sporozoite numbers and motility were overall similar to those in wild type infections (Figure 3A-C). The *P. falciparum* orthologue is thus functional in *P. berghei* even though it has a long N-terminal extension and an overall identity of only 35% (see Figure S2).

**Fig. 3.**
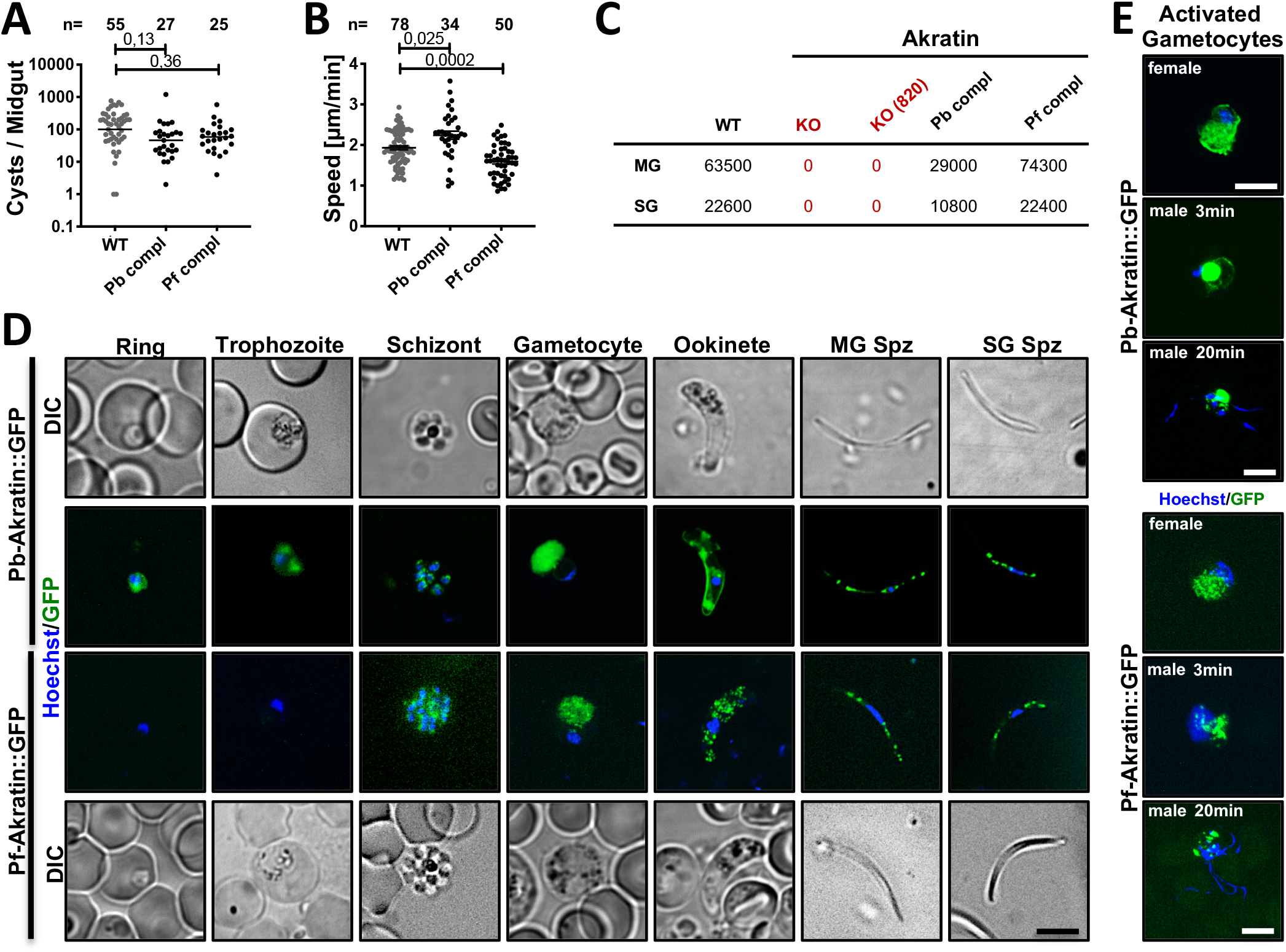
Complementation of the *akratin(-)* parasites with akratin-GFP restores the wild type phenotype and shows stage dependent cytoplasmic and vesicular localization. **(*A*)** Oocyst development of *akratin(-)* parasites complemented with either the *P. berghei* gene or the *P. falciparum* orthologue fused to GFP. Data points represent individual midguts observed between d12-17 post infection. Horizontal line shows median. P-values are calculated using the Kruskal Wallis test followed by the Dunns multiple comparison test. **(*B*)** Speed of salivary gland sporozoites imaged with a frame rate of 3 seconds. Data points represent average speed of individual sporozoites. Shown is the mean ± SEM. P-values calculated by Kruskal Wallis test followed by Dunns multiple comparison test. (***C)*** Sporozoite numbers in midgut (MG) and salivary glands (SG) from infections by wild type, *akratin(-), P. berghei* and *P. falciparum* complementation counted on d17 post mosquito infection. For each counting at least 20 mosquitos were dissected. **(*D*)** *Pb*akratin::gfp and *Pf*akratin::gfp localization in blood and mosquito stages. Nuclei (blue) are stained with Hoechst. DIC: differential interference contrast, MG: midgut, SG: salivary gland. Note the weaker signal in *P. falciparum* akratin-GFP in gametocytes and the difference between the signals in ookinetes. Scale bar: 5μm. **(*E*)** akratin::gfp localization in activated male and female gametocytes. Scale bar: 5μm.

We next investigated the localization of the fluorescent signal in the blood stage and the extracellular forms of the life cycle. This revealed a mostly diffuse signal with some punctae in blood stages of both parasite lines expressing the *P. falciparum* or *P. berghei* akratin-GFP (Figure 3D). In non-activated gametocytes the signal was distributed within the gametocyte cytoplasm but it was much weaker in the *P. falciparum* akratin-GFP line, where it appeared punctate. No difference between male and female gametocytes could be observed. However, upon activation two distinct populations, most likely males and females could be observed. While in females the protein was distributed in the cytoplasm of the cell, male (determined by a cloudy Hoechst staining) showed one bright spot in addition to a few weak dots at the periphery (Figure 3E). Ookinetes showed a vesicular/endomembranous staining with a prominent peripheral staining for the *P. berghei* akratin-GFP but an exclusively vesicular pattern for the *P. falciparum* akratin-GFP (Figure 3E; Figure S7). Sporozoites isolated from the oocysts or salivary glands showed a similar punctate localization for both lines reminiscent of secretory vesicles but distinct from the strong apical signal seen in GFP-TRAP sporozoites (Kehrer, Singer, et al., 2016).

### Akratin is essential for microgametogenesis

To probe if akratin functions in male or female gametocytes we performed parasite-crossing experiments with the *akratin(-)* parasite line and a parasite line that either produces only fertile males, *p47(-)*, or only fertile females, *p48/45(-)* (van Dijk et al., 2010). As positive control we crossed the *p47(-)* and *p48/45(-)* lines with each other. This control as well as the *akratin(-)* x *p47(-)* cross resulted in ookinete conversion rates of 35% and 42%, respectively, while the *akratin(-)* x *p48/45(-)* cross produced only few ookinetes (3%) (Figure 4A), suggesting that female *akratin(-)* gametocytes are still fully functional, while male gametocytes are strongly impaired.

**Fig. 4.**
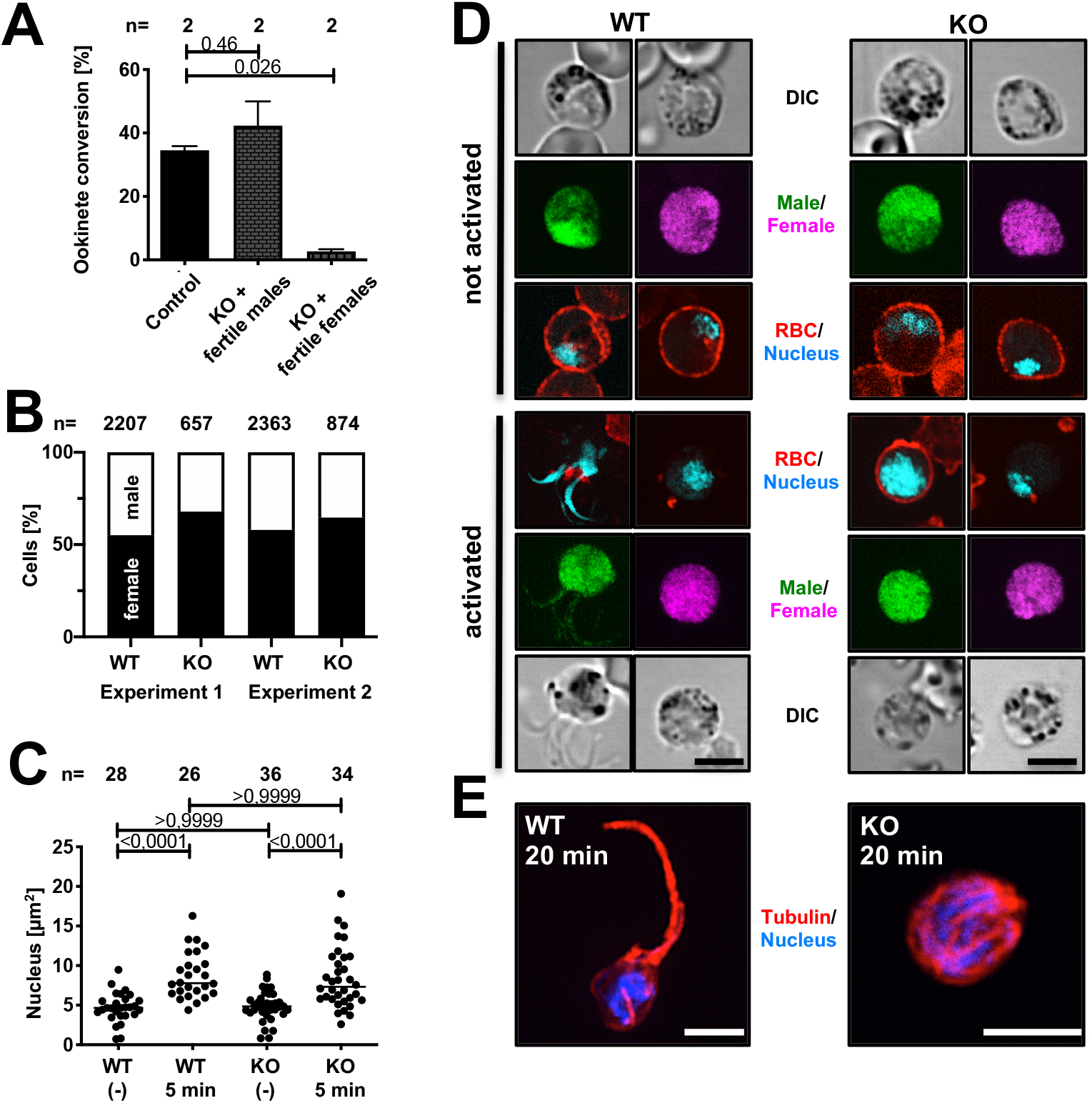
*Akratin(-)* parasites do not complete microgametogenesis. **(*A*)** Crossing of parasite lines deficient in either male or female gametocytes with *akratin(-)* parasite reveals that akratin functions only in males. P-value calculated by One-way Anova with Dunnetts multiple comparison test. Shown is the mean±SEM**. (*B*)** The same number of male and female gametocytes are formed in wild type (WT) and *akratin(-)* (KO) parasites lines. **(*C*)** Nuclear size determined from non-activated (-) or activated (5 min) wild type or *akratin(-)* parasites. Data points represent size of individual nuclei. P-values calculated by Kruskal Wallis Test with Dunns multiple comparison. (***D*)** Activation of gametogenesis releases male and female WT gametes but only female *akratin(-)* gametes. Red blood cell (RBC) membrane is stained with Ter119 (red); male parasites are shown in green, females in pink and DNA in cyan. Scale bar: 5μm. **(*E*)** 20 minutes post activation *akratin(-)* parasites show microtubule staining reminiscent of assembled axonemes but not egressed gametes, while wild type parasites show the typical axonemal staining of exflagellating gametes. Scale bars: 5 μm.

To investigate if akratin functions in male gametocytogenesis we generated an additional *akratin(-)* parasite line in the 820cl1m1cl1 (RMgm-164) background (Figure S5). These selection marker-free parasites express RFP in females and GFP in male gametocytes (Ponzi et al., 2009). The ratio between both sexes in the resulting *akratin(-)-820* line did not show any difference between the wild type and the KO (Figure 4B) suggesting that male gametocytes are still normally formed. To investigate at which stage the cells arrest during microgamete formation, we activated the gametocytes and fixed them at different time points. Male gametogenesis involves 3 rounds of rapid DNA replication, which is accompanied by an increase of the size of the nucleus within the first minutes (C. J. Janse et al., 1986). DNA staining with Hoechst of activated and non-activated cells showed no difference in nuclear size between *akratin(-)* and wild type parasites (Figure 4C), indicating that *akratin(-)* microgametes are activated and replicate their DNA normally. Staining of the red blood cell membrane, however, revealed that it is not degraded in male *akratin(-)* parasites (Figure 4D) providing a possibly function for akratin in membrane lysis. Supporting such a function, anti-tubulin antibodies revealed normally formed flagellar axonemes albeit non motile, that appeared trapped within the red cell in *akratin(-)* gametocytes, while wild type gametes readily extended from the cell (Figure 4E).

### Mutation of a C-terminal motif uncouples akratin function in gametocytes and ookinetes

In mammalian cells the C-terminal sorting sequence YXXφ (with Y, tyrosine; X, any amino acid and φ, hydrophobic amino acid) is important for protein trafficking (Marks et al., 1997). A similar motif was also shown to be important for trafficking of the micronemal sporozoite protein TRAP (Bhanot et al., 2003), although the sequence is annotated as being located in the trans-membrane domain (Figure 5A). Somewhat similar tyrosine-containing C-terminal motifs (SYHYY and EIEYE) from the *Toxoplasma gondii* adhesin MIC2 were also shown to target the protein to micronemes (Di Cristina et al., 2000). We noticed in the C-terminal, likely cytoplasmic domain of *P. berghei* akratin a short sequence (YKKL) representing the described YXXφ pattern (Figure 5A). However, since the *P. falciparum* orthologue contains a leucine instead of a tyrosine and a conserved glutamic acid to the left we chose to mutate the sequence EYKK instead of YKKL to investigate a possible role for akratin trafficking. To this end, we generated a mutant parasite line expressing akratin-GFP with the EYKK motif changed into four alanines (Figure S6). Akratin-^EYKK/AAAA^-GFP showed similar staining patterns in asexual and sexual parasite stages compared with akratin-GFP (Figure 5B). However, in contrast to the mostly peripheral and vesicular localization of akratin-GFP in ookinetes the akratin-^EYKK/AAAA^-GFP signal was diffusely localized within the cytoplasm (Figure 5B; compare with Figure 3F; Figure S7). Although the exflagellation was somewhat reduced, the mutant did readily form ookinetes at similar rates as wild type and was able to move in a wild type manner (Figure 5C-E). Strikingly, however, despite their capacity to migrate on glass, the mutant ookinetes failed to form oocysts (Figure 5F). To investigate if this defect is due to the failure to penetrate the mosquito midgut or due to a failure in oocyst formation, we isolated midguts 24h hours post infection, washed and stained them for the actin cytoskeleton outlining the epithelial cells and the muscles fibers on the hemolymph-facing side of the midgut (Figure 5G) and searched for ookinetes. This showed half the investigated wild type infected midguts with ookinetes that penetrated the epithelial layer. However, none was found in the mutant (Figure 5G), suggesting that akratin is differentially trafficked in gametocytes and ookinetes and that akratin is needed on the ookinete surface for interaction with epithelial cells or their traversal. Parasites lacking akratin appear powerless to enter the epithelium, a second reason for their name.

**Fig. 5.**
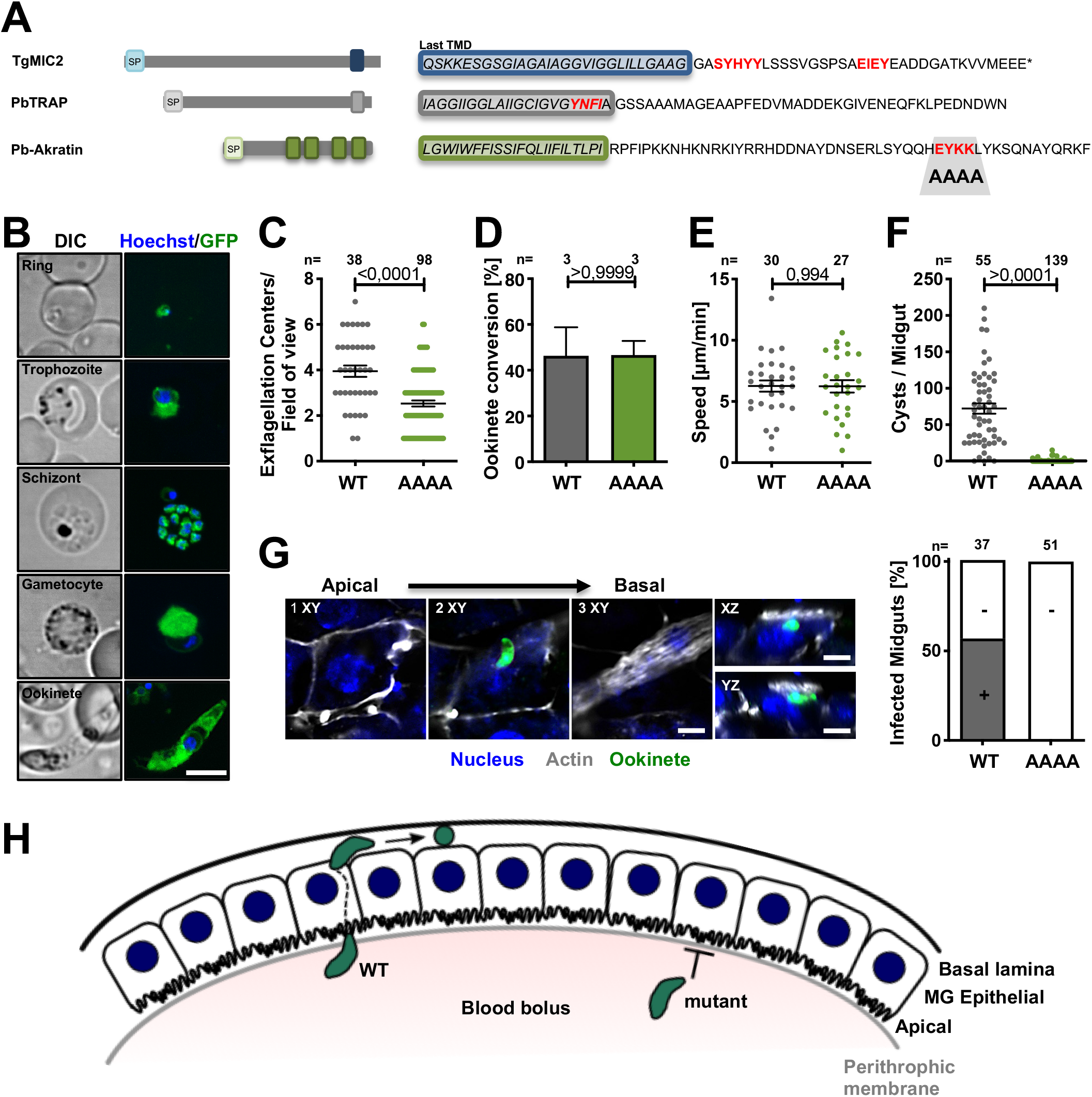
Mutation of a putative trafficking motif mislocalizes akratin-GFP in ookinetes and ablates oocyst formation. **(*A*)** Cartoon showing signal peptides (SP) and trans-membrane domains (TMD) of the *T. gondii* adhesin MIC2 and *P. berghei* Trap and akratin. C-terminal sequences with its putative micronemal targeting motifs are highlighted in red. **(*B*)** Localization of akratin::GFP in the indicated *P. berghei* life cycle stages. Scale bar: 5μm. **(*C*)** Exflagellation is almost normal in the akratin^EYKK/AAAA^ mutant. P-value calculated by Mann Whitney test. Shown is the mean±SEM. **(*D*-*E*)** Ookinete conversion rate is not affected in the EYKK/AAAA mutant **(*D*)** and motility **(*E*)** is comparable to wild type (WT). P-value calculated by Mann Whitney test. Shown is the mean±SEM. **(*F*)** Oocyst numbers a significantly reduced in akratin^EYKK/AAAA^ parasites. P-value calculated by Mann Whitney test. Shown is the mean±SEM. **(*G*)** Akratin^EYKK/AAAA^ ookinetes fail to penetrate the mosquito midgut epithelium while more than half of WT midguts showed ookinetes. Confocal images in the indicated views shows wild type ookinetes at different positions within the epithelial layer. Scale bar: 5μm. Graph shows the number of infected and not infected midgut epithelia; no akratin^EYKK/AAAA^ mutants were found in the epithelial cell layer or beyond. **(H)** Cartoon: the path of wild type (WT) and mutant ookinetes from the blood meal across the epithelial layer. The akratin^EYKK/AAAA^ mutant arrests below the basal membrane.

## Discussion

Using APEX2-based rapid proximity labeling for the first time in Apicomplexa, we defined the content of secretory vesicles in *Plasmodium berghei* ookinetes. With just two experiments we successfully identified all well-known micronemal ookinete proteins as well as over 50 hitherto unknown candidates, some of which can also be found in the sporozoite proteome and might thus also be important for *Plasmodium* transmission from the mosquito to the mammal. Classifying these candidates into proteins with signal peptides and transmembrane regions allowed us to select a candidate, named akratin, for functional analysis. Akratin-GFP localized in all investigated stages of the life cycle with different patterns, appearing cytosolic in both asexual and sexual blood stages and more vesicular in ookinetes and sporozoites. Deletion of *akratin* resulted in a complete block of parasite transmission into mosquitoes but surprisingly showed a phenotype already in the exflagellation of male gametes nearly abolishing gamete release and hence ookinete formation. Other membrane proteins such as PAT (Kehrer, Singer, et al., 2016) or MTRAP (Bargieri et al., 2016; Kehrer, Frischknecht, et al., 2016) have also been shown to play a role in gamete exit from red blood cells, although their detailed mode of action is not clear.

Akratin also showed an important function in ookinetes, revealed by the mutation of a putative microneme targeting motif. This mutation had no detrimental effect on akratin function during gamete formation, where akratin was localized within the cytoplasm rather than within vesicles, an unexpected finding for a protein containing four putative trans-membrane domains. In ookinetes mutation of the motif reduced the targeting to the plasma membrane. Ookinetes still formed normally and were also equally motile as wild type control parasites on glass, yet they failed to penetrate the midgut epithelium and thus could not develop into oocysts. How can ookinetes still move *in vitro* but not migrate *in vivo?* One possibility is that they fail to bind a key receptor and hence do not signal for a potential switch from migration to invasion. Or the mutant ookinetes do not produce enough force and thus cannot penetrate through the peritrophic matrix that builds around the digesting blood meal or enter the epithelium. We recently observed a similar effect when replacing the essential ookinete adhesin CTRP with the sporozoite adhesin TRAP (Klug et al., 2018). Without CTRP ookinetes do not glide on glass and also do not enter the epithelium. Strikingly, the TRAP expressing ookinetes moved on glass just fine but did not form oocysts, suggesting that *in vitro* motility is not a perfect predictor for *in vivo* migration. Indeed, several mutants generated in our laboratory show a similar disconnect for sporozoites and their capability to enter into salivary glands. Some show a strong defect in gliding on glass, yet enter normally into salivary glands (Moreau et al., 2017), while others migrate almost normally but do not enter efficiently into glands (Douglas et al., 2018). These mutants show mutations in actin or actin-binding proteins and hence should not affect receptor-ligand interactions, although we speculate that receptor-ligand interactions might cause a rearrangement of actin, which could feed back to modulate the receptor-ligand binding strength (Quadt et al., 2016). A similar complex outside-in to inside-out signaling might well play a part in the complex journey of the ookinete, which needs to first escape the blood meal, cross the peritrophic matrix, migrate along and through the epithelial cell layer and arrest below the basal lamina, a journey certainly benefiting from fine-tuned sensing of environment. Another possibility for the observed defect of *akratin(-)* penetration of the epithelium would be that the protein protects the ookinete from the onslaught of the vertebrate or mosquito defense proteins such as TEP1 (Fraiture et al., 2009).

In conclusion, we established a new assay for rapid biotinylation in malaria research, generated a list of putative micronemal proteins in ookinetes, which might harbor new transmission blocking candidates and identified a protein, akratin, with dual roles in gamete formation and ookinete migration. We furthermore showed that a putative micronemal targeting motif is important for proper targeting and function of akratin in ookinetes but not in gametocytes. While the discovery of akratin opens interesting biological questions the use of rapid BioID will provide an important tool for discovery along the complex *Plasmodium* life cycle.

## Materials and Methods

### Generation of transfection plasmids

#### soap::APEX2

The *APEX2::myc* sequence flanked with the restriction sites BamHI and XbaI was ordered from geneart (Regensburg, Germany). It was ligated into the previously described BioID plasmid (Kehrer, Frischknecht, et al., 2016) replacing the *bira\**-sequence. The resulting plasmid pL8 was linearized with HpaI prior transfection for single crossover integration (Figure S1).

#### akratin(-)

The 3’UTR of PbANKA_1105300 was amplified using primers JK67 and JK68 and inserted into a plasmid containing the recyclable yFCU/ hDHFR selection cassette and GFP expressed under the HSP70 promoter digested with NotI and SacII. The 5’UTR of PbANKA_1105300 was amplified using primers JK65 and JK66 and inserted into the plasmid using KPNI and HindIII. The resulting plasmid pL28 was linearized with NotI and SacII prior transfection for double crossover integration (Figure S3).

#### Marker free *akratin(-)*

The drinking water of mice was supplemented with 2 mg/ml 5-FC. Clonal parasites which looped out the selection cassette, were obtained by limiting dilution (Figure S3).

#### *P. berghei akratin::gfp* complementation

The 5’UTR together with the entire ORF of PbANKA_1105300 was amplified from wt gDNA using primers JK66 and JK152 and inserted into the pL28 plasmid using KpnI and NdeI leading to the replacement of the selection marker. A TgDHFR selection cassette was amplified using primers JK153 and JK154 and inserted between the GFP and 3’UTR using NotI and EcorV resulting in plasmid pL59. For transfection of parasites via double crossover the plasmid was digested with KpnI and SacII (Figure S4).

#### *P. falciparum akratin::gfp* complementation

The ORF of PF3D7_0505700 was amplified using primers JK171 and JK172 and inserted into pL28 downstream of the *P. berghei* 5’UTR using HindIII and NdeI leading to the replacement of the selection marker. The TgDHFR selection cassette was amplified using primers JK153 and JK154 and inserted between the GFP and 3’UTR using NotI and EcorV resulting in plasmid pL75. For transfection of parasites via double crossover the plasmid was digested with KpnI and SacII (Figure S4). *akratin^EYKY/AAAA^:* The 5’UTR together with the entire ORF was amplified from wt gDNA using primers JK66 and a reverse Primer introducing the mutations and inserted into the pL59 plasmid using KpnI and NdeI resulting in pL84. For transfection of parasites via double crossover the plasmid was digested with KpnI and SacII and transfected into negative selected KO parasites (Figure S6).

### Generation of transgenic parasite lines

Transfection of linearized plasmids was performed as previously described (Chris J. Janse et al., 2006). After electroporation of schizonts, positive selection was performed using pyrimethamine. All resulting lines were cloned through limiting dilution except *soap::APEX2* and *akratin(-). soap::APEX2* parasites still contain a small amount of wild type parasites that did not interfere with BioID, while the *akratin(-)* was obtained through FACS sorting as follows: 2 drops of blood from the mouse tail vein with a parasitemia of maximum 0.3 % were diluted in 1.5 ml of RPMI medium. To avoid clustering of cells the sample was drained before analysis, with a 40 μm cell strainer. For detection, cells were analysed with a 488 nm laser in combination with a 527/32 nm filter for the GFP signal. To set up gates for sorting, the voltage of forward scatter (FSC) and sideward scatter (SSC) were adjusted to mainly gate on erythrocytes and a singlet discrimination was performed. Cells were sorted in the purity mode with the lowest flow rate at room temperature using a BD FACS Melody. As sheath fluid PBS was used. Sorted cells (hundred events) were immediately injected i.v. into naïve mice. No wild type could be detected in the resulting polyclonal *akratin(-)* population.

### *In vitro* ookinete cultures and proximity labeling

20 million blood stage parasites were injected i.p. into a naïve NMRI mouse treated with 100μl phenylhydrazin (6μl/ 1ml) 24h pre-transfer. 3 days post infection the drinking water was supplemented with sulfadiazin (20 mg/L) to reduce asexual blood stages. 4 days post infection the blood was harvested by cardiac puncture to set up ookinete cultures. The blood of one mouse was added to 10 ml of ookinete medium (RPMI containing 25 mM Hepes and 300 mg l^−1^, l-glutamine, 10 mg l^−1^ hypoxanthine, 50 000 units l^−1^ penicillin, 50 mg l^−1^ streptomycin, 2 g l^−1^ NaHCO_3_, 20.48 mg l^−1^xanthurenic acid, 20% FCS; pH 7.8) at 19°C and incubated for 21 hours. Fully developed ookinetes were purified on a 63% Nycodenz cushion and the remaining RBCs were lysed with 170 mM NH_4_CL for 5min on ice.

Cells were washed once with PBS and resuspended in 1 ml of ookinete medium containing biotin phenol (182 μg/ml). After incubation for 30 min at RT cells were split into 2 tubes. One was used as control and in one biotinylation was activated by adding 5μl of 100mM H_2_O_2_. After 90 seconds the reaction was inactivated by adding 500 μl of 2x quencher solution (20 mM Sodiumascorbate, 20 mM Trolox, 10 mM NaN3 in PBS). After washing the cells two times with 500 μl 1x quencher solution samples were then lysed with RIPA buffer (50 mM Tris pH8, 1% Nonidet P-40, 0.5% Na-deoxycholate, 0.1% SDS, 150 mM NaCl, 2 mM EDTA). 20% of the lysate was kept as input control and biotinylated proteins were enriched using streptavidin coated beads at 4°C overnight as described (Roux et al., 2012). Elution of proteins was done with a buffer containing 30 mM biotin, 2% SDS, 160 mM NaCl and 6 M Urea followed by mass-spectrometric analysis as described below.

### Protein analysis and proteomics

Western blots were performed using the following antibodies: anti-myc antibody (Roche 0.4 mg/ml, 1/1000) followed by incubation with goat anti mouse HRP (BioRad 1/10000), Streptavidin-HRP (Sigma, 1/500) or mouse anti-HSP70 followed by incubation with goat anti-mouse HRP. Mass spectrometric analysis was performed at the CellNetworks Core Facility for Mass Spectrometry and Proteomics of the ZMBH (Zentrum für Molekularbiologie der Universität Heidelberg) as previously described (Kehrer, Frischknecht, et al., 2016).

### Immunofluorescence and light microscopy

After fixation of the parasites with 4% PFA at 4°C for at least overnight, cells were permeabilized with 0.5% TritonX-100 for 10 min. Staining of the red blood cell membrane was performed with the anti-Ter119::Alexa647 antibody (Biolegend, 0.5 mg/ml; 1/1000) for 1 hour. Cells were washed twice with PBS, and resuspended in PBS containing Hoechst for observation under the microscope.

For staining of microtubules cells were fixed in 2% Paraformaldehyde/0.05% Glutaraldehyde in microtubule stabilizing buffer (15 g/L PIPES, 1.9 g/L EGTA, 1.32 g/L MgSO_4_·7H_2_O and 5 g/L KOH in water; pH 7.0 with KOH). Staining was performed using an a-tubulin antibody (Sigma clone DM1A, 1/ 500) for 1 hour, followed by an anti-mouse 647 (1/ 500) for 1 hour. Images were either taken on a Zeiss 200 M Axio-Observer widefield (63x; NA 1,4) or Nikon/PerkinElmer spinning disc confocal (100x; NA 1,4) microscope. Image processing was performed with ImageJ/FIJI (Schindelin et al., 2012).

### Ookinete motility and midgut traversal

*In vitro* ookinete cultures were performed as described above. To observe motile ookinetes one drop of cell suspension was transferred onto a microscope slide and covered with a thin cover glass to simulate a confined environment. Movies were taken with a frame rate of 20 seconds for 10 minutes. Speed was analyzed using the manual tracking plugin in FIJI.

For *in vivo* analysis, 20 million blood stage parasites were injected i.p. into a naive Swiss CD1 mouse treated with phenylhydrazin 24 hours prior to transfer. 4 days post transfer about 20 female *Anopheles* mosquitoes were allowed to feed for about 15 minutes. Blood filled midguts were isolated after 24 hours as described previously (Han et al., 2000). Midguts were fixed in cold 4% paraformaldehyde (PFA) for 45 seconds and washed with cold PBS. The epithelial cell layer was then opened longitudinally using two needles to carefully remove the blood bolus followed by a second fixation with 4% PFA overnight.

To better visualize traversed ookinetes an additional immunofluorescence assay was performed. To do so cells were permeabilized with 0,5% Triton X-100 for 30 minutes, incubated with a rabbit anti GFP antibody (abfinity, 0,4μg/μl, 1/40) for 2 hours and after washing incubated with an anti rabbit 488 secondary antibody plus the addition of Phalloidin TRITC (1mg/ml, 1/500) and Hoechst (10mg/ml; 1/1000) for another 2 hours. Midguts were transferred onto a microscope slide and images were taken on a Nikon spinning disc microscope using either a 40x (N.A. 0,65) or 60x (N.A. 1,4) objective.

## Acknowledgements

We thank Miriam Reinig for *Anopheles stephensi* mosquito rearing, Gunnar Mair for help with initial experiments on SOAP-BirA*-tagging and fruitful discussions, Blandine Franke-Fayard and Chris Janse for the 820 line and Markus Ganter for comments on the manuscript. Mass spectrometry and proteomics analysis was performed at the *CellNetworks-ZMBH* core facility at Heidelberg University. We especially thank Bernd Hessling for his support. We also thank Carolina Barillas-Mury for helpful discussions on ookinete midgut traversal. We acknowledge support from the Infectious Diseases Imaging Platform (IDIP) at the Center for Integrative Infectious Disease Research. Funded by the Deutsche Forschungsgemeinschaft (DFG, German Research Foundation) – Project-number 240245660 - SFB 1129, Human Frontier Science Program Young Investigator grant RGY066 and the European Research Council (StG 281719). FF is a member of the *CellNetworks* Cluster of Excellence (EXC 81) at Heidelberg University. DR, LS, EP and JH were members of the master programs of Molecular Biotechnology (DR) or Infectious Diseases at Heidelberg University.

## Footnotes

To whom correspondence may be addressed.

Author contributions: J.K. and F.F. designed research; J.K., D.R., L.S., E.P. and J.H. performed research; all authors analyzed data; and J.K. and F.F. wrote the paper.

The authors declare no conflict of interest

This article contains supporting information online

## Supplementary Figures

**Fig. S1.**
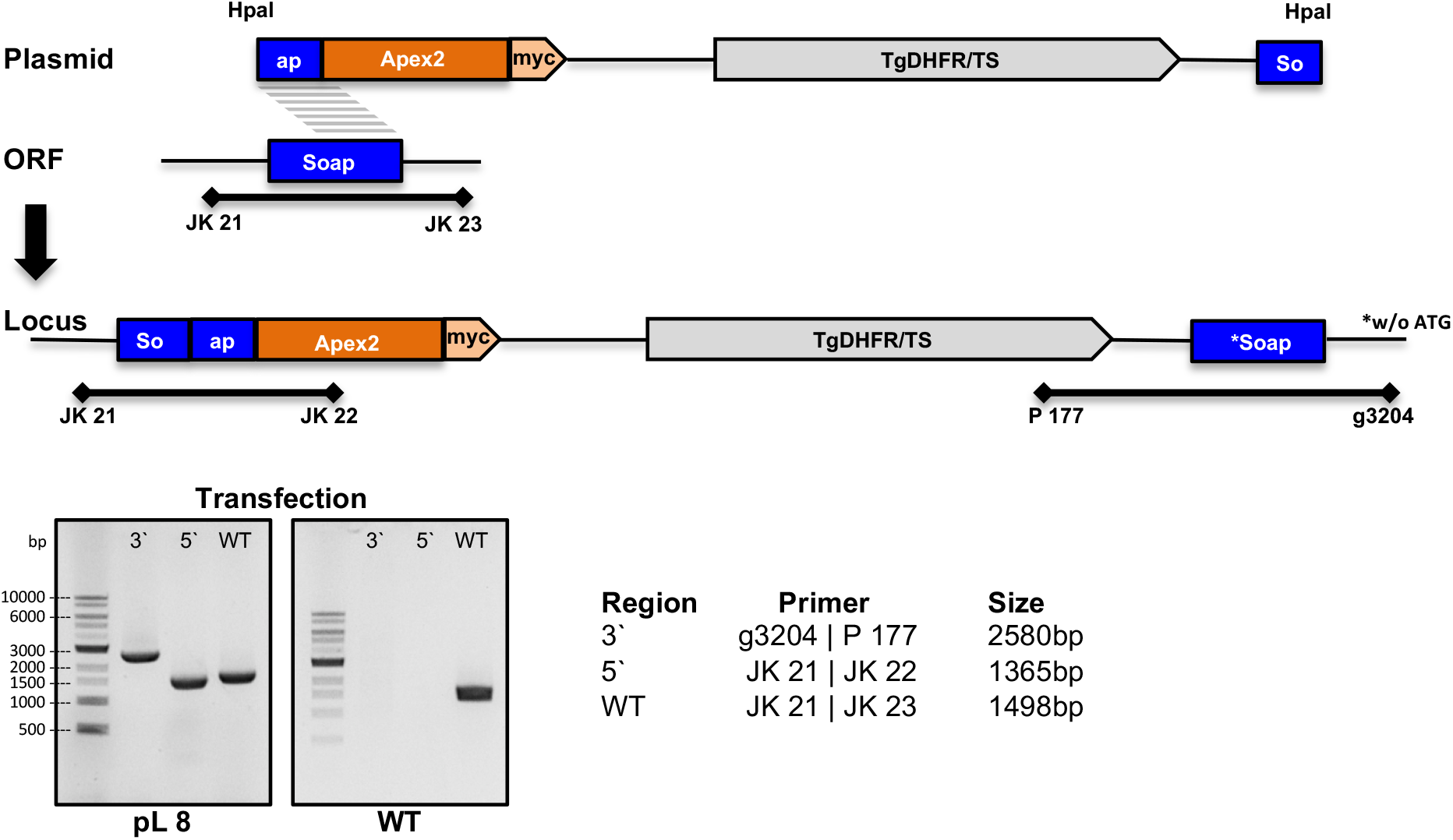
Generation of Soap::APEX2 parasites via single homologous recombination. The cartoon shows the cloning strategy and primers used for genotyping with amplicon sizes of the resulting transgenic line indicated. For primer sequences please see table S2. Plasmid contains as resistance marker the dehydrofolatereductase/thymidine-synthase from *Toxoplasma gondii* (*Tgdhfr/ts*) (grey). Note that the second copy of soap lacks the ATG and should not be expressed.

**Fig. S2.**
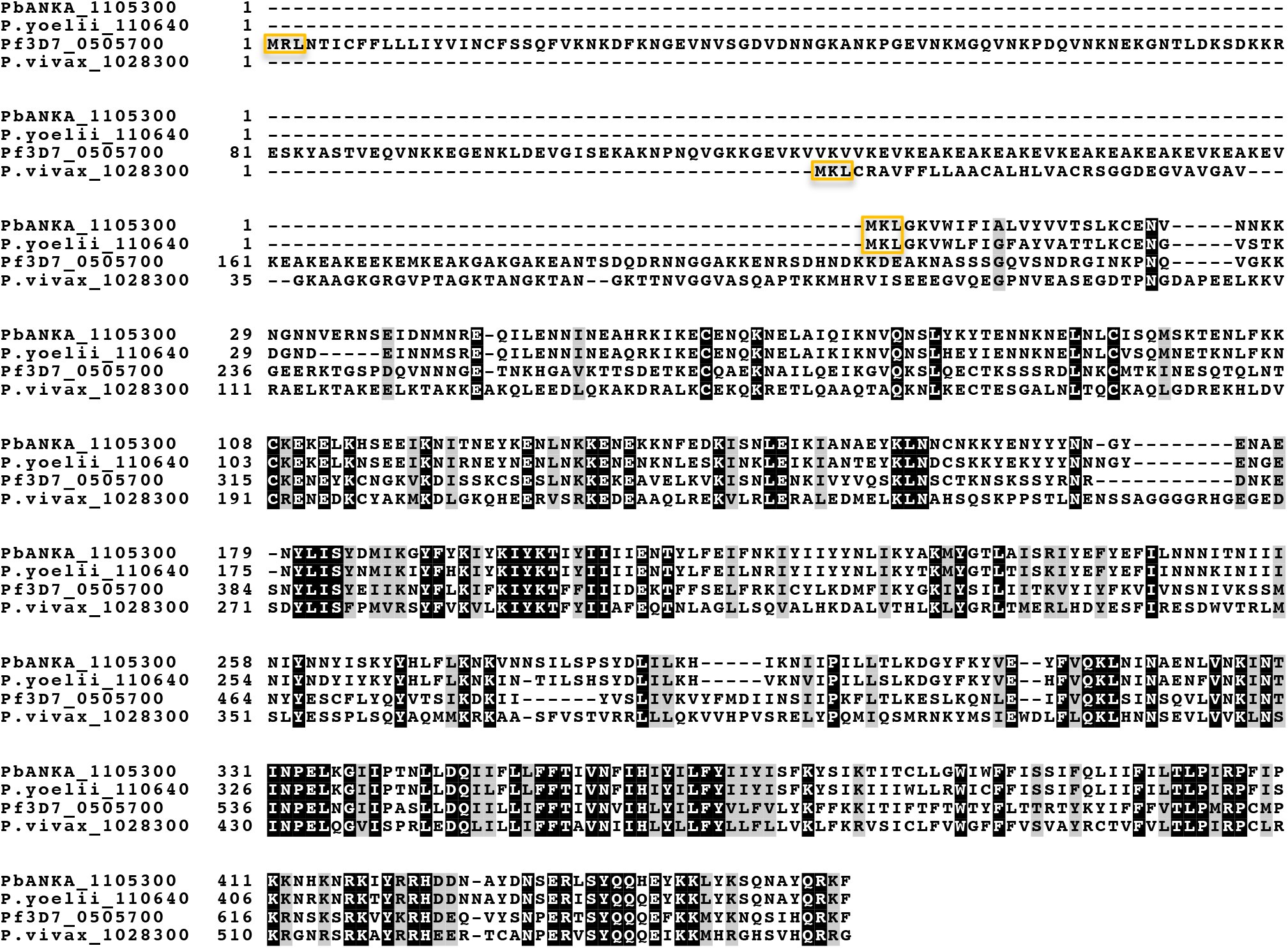
Multiple protein sequence comparison between *Plasmodium spp*. using Clustal Omega alignment. Highlighted are the bases conserved in all species (black) and respective start amino acids (orange).

**Fig. S3.**
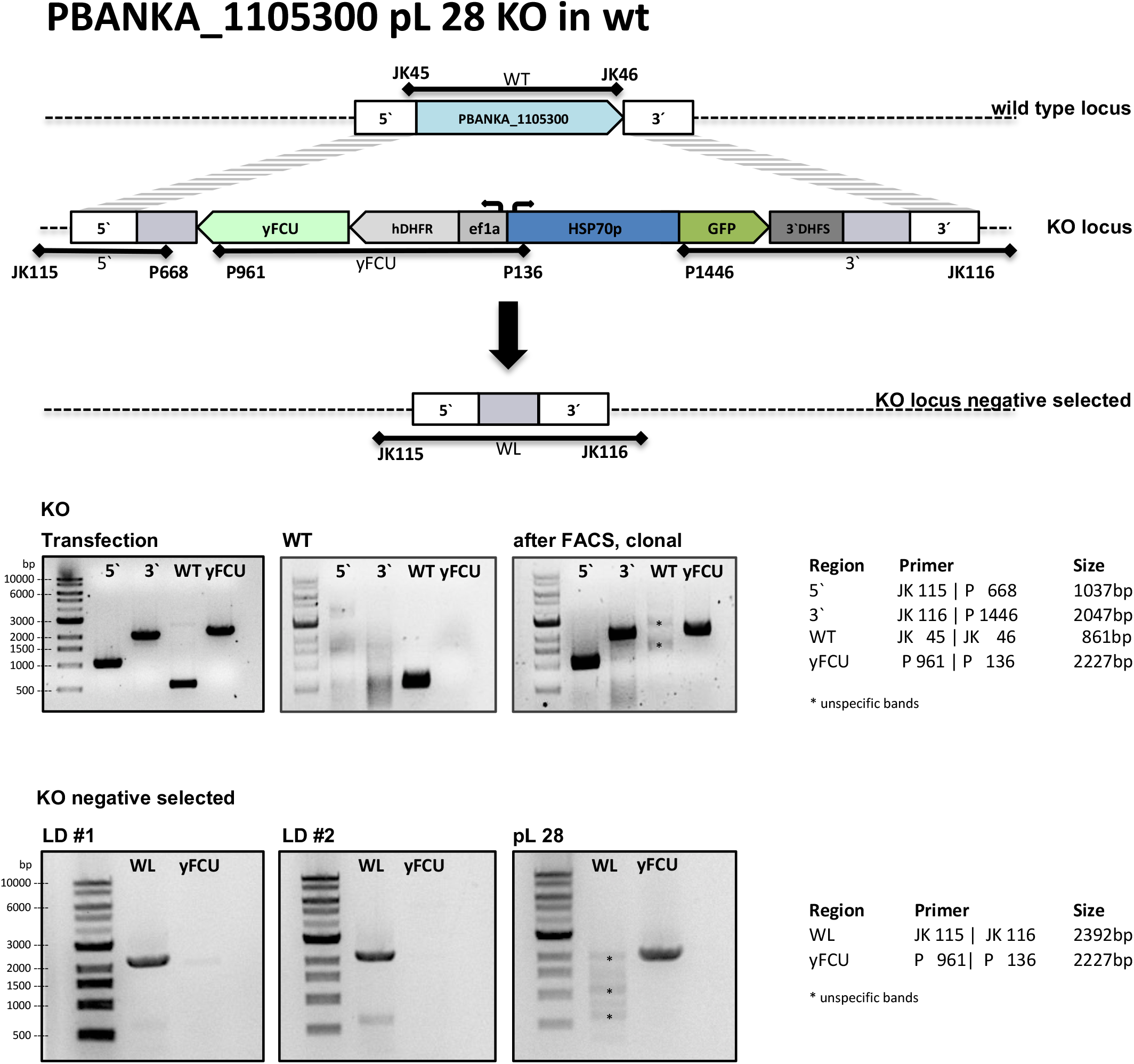
Generation of *akratin(-)* and *akratin(-)* negative selected parasites via double homologous recombination. The cartoon shows the cloning strategy and primers used for genotyping. Note that as yFCU (Yeast cytosine deaminase-uracil phosphoribosyl transferase fusion protein) was used as a negative selection marker (see methods).

**Fig. S4.**
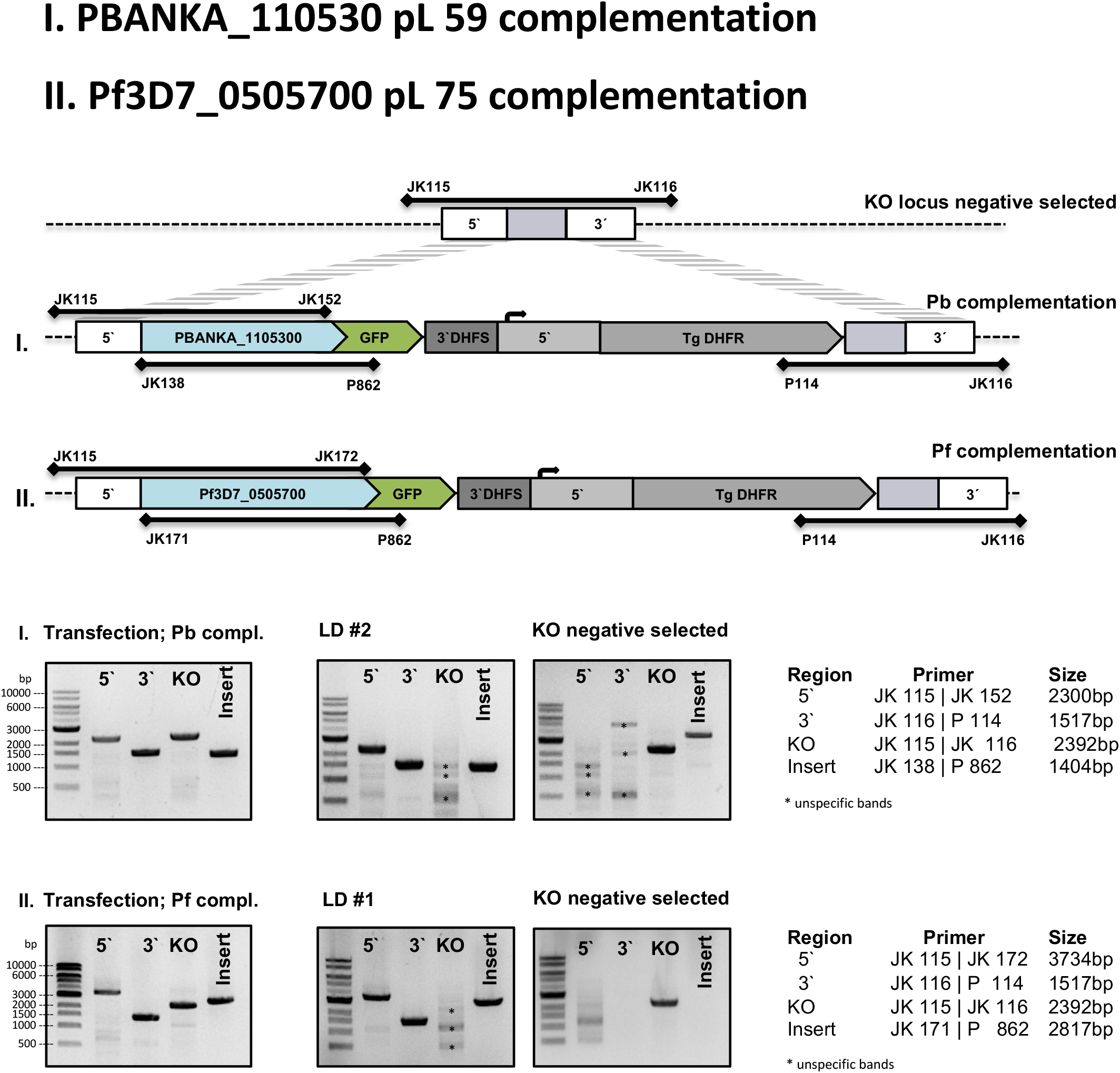
Generation of *P. berghei* and *P. falciparum* complementation parasites via double homologous recombination. The cartoon shows the cloning strategy and primers used for genotyping.

**Fig. S5.**
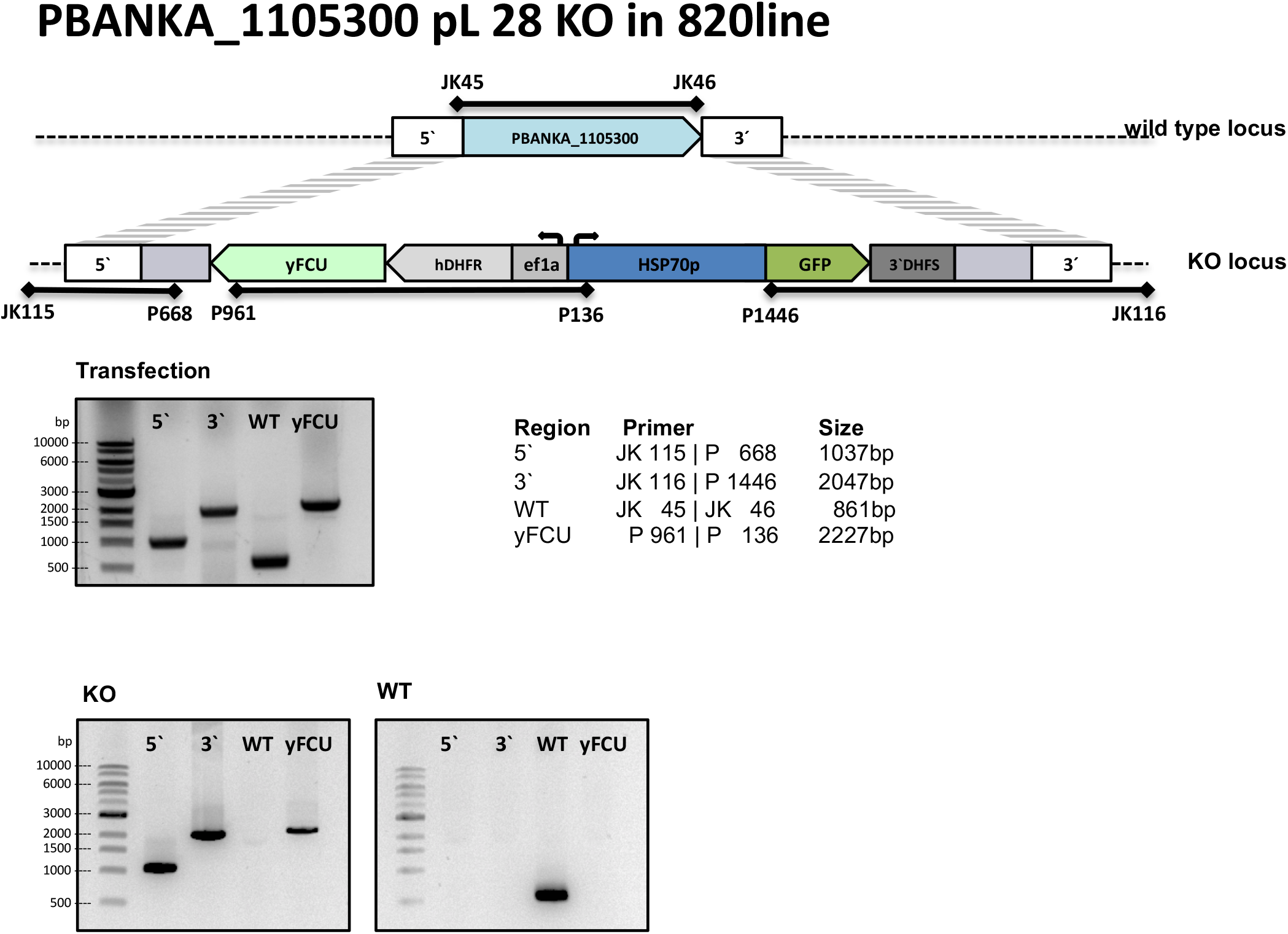
Generation of *akratin(-)* parasites in the 820line via double homologous recombination. The cartoon shows the cloning strategy and primers used for genotyping.

**Fig. S6.**
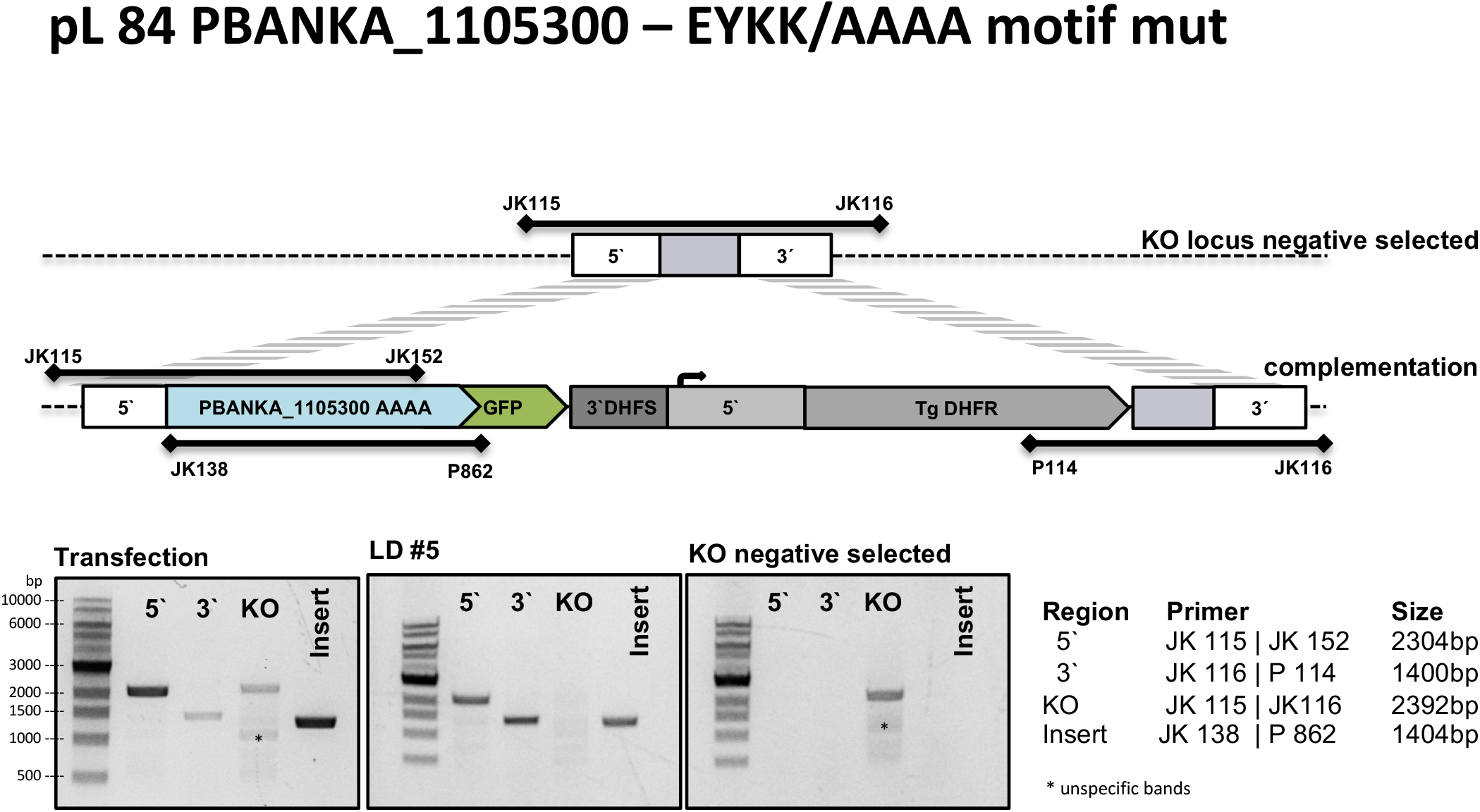
Generation of akratin^EYKK/AAAA^ parasites via double homologous recombination. The cartoon shows the cloning strategy and primers used for genotyping.

**Fig. S7.**
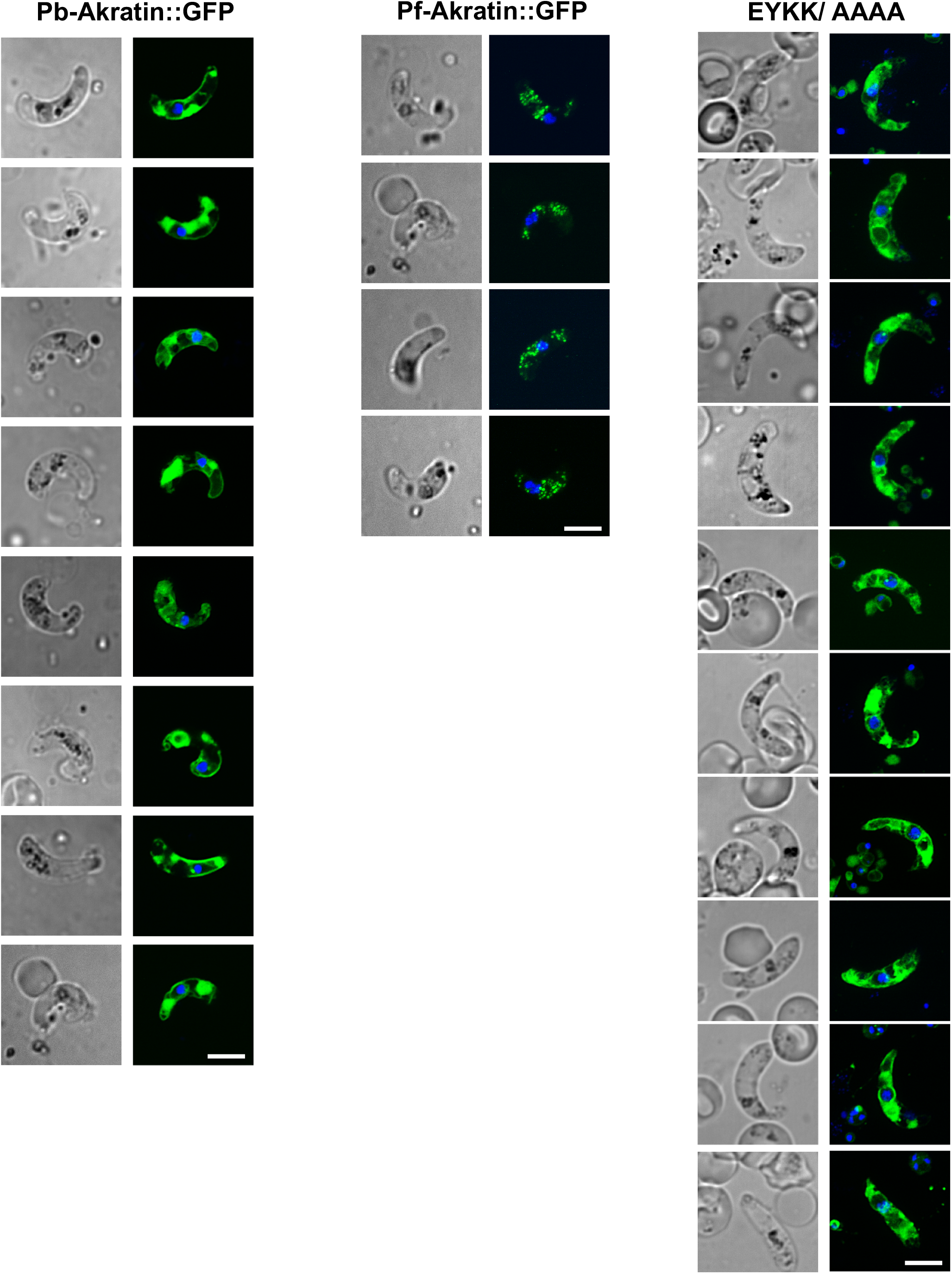
Gallery of ookinetes expressing PbANKA_1105300::GFP, PF3D7_0505700::GFP or EYKK/AAAA::GFP. Scale bar 5μm.

**Table S1.**
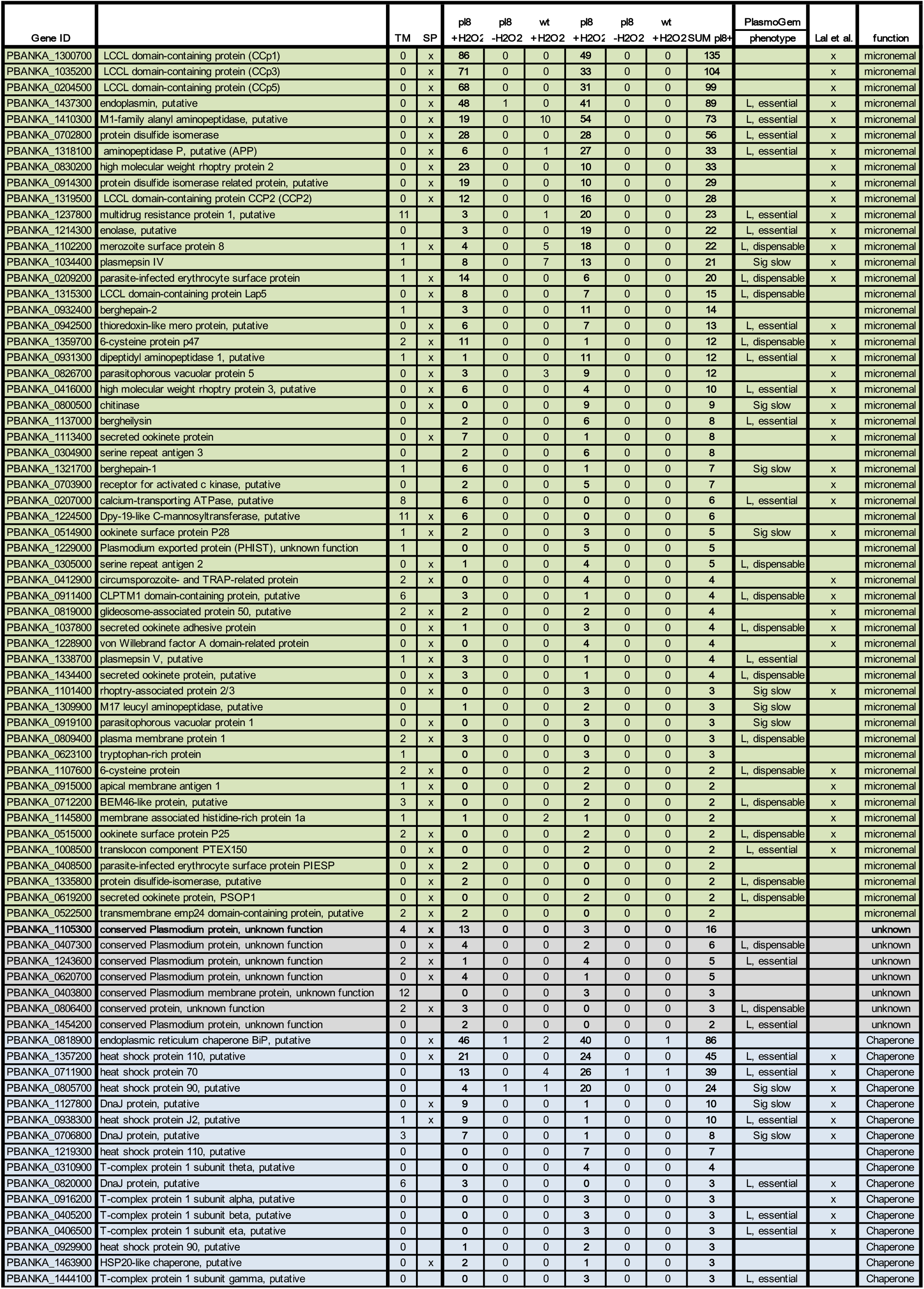
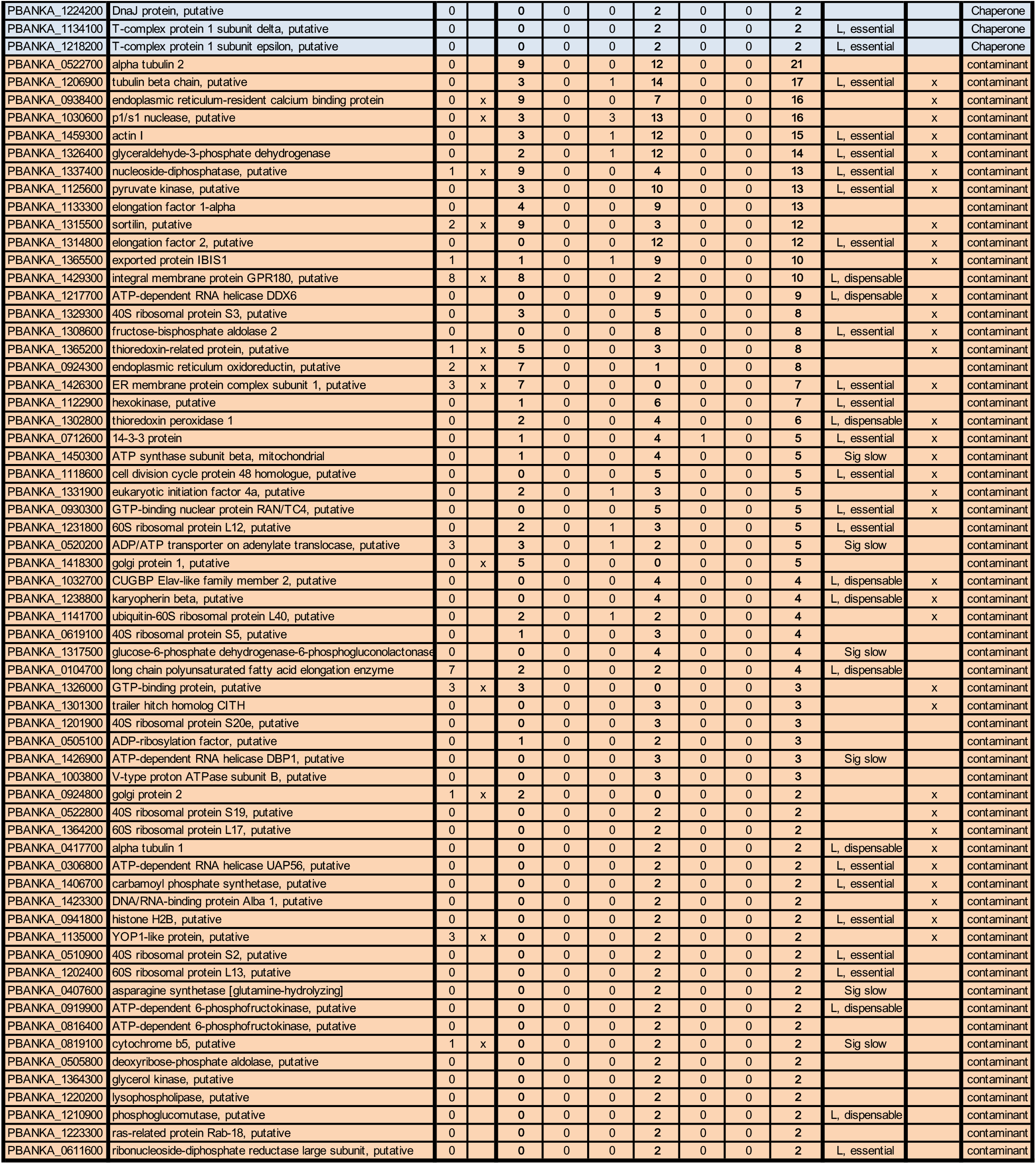
Proteomic analyses of ookinete micronemes

**Table S2.**
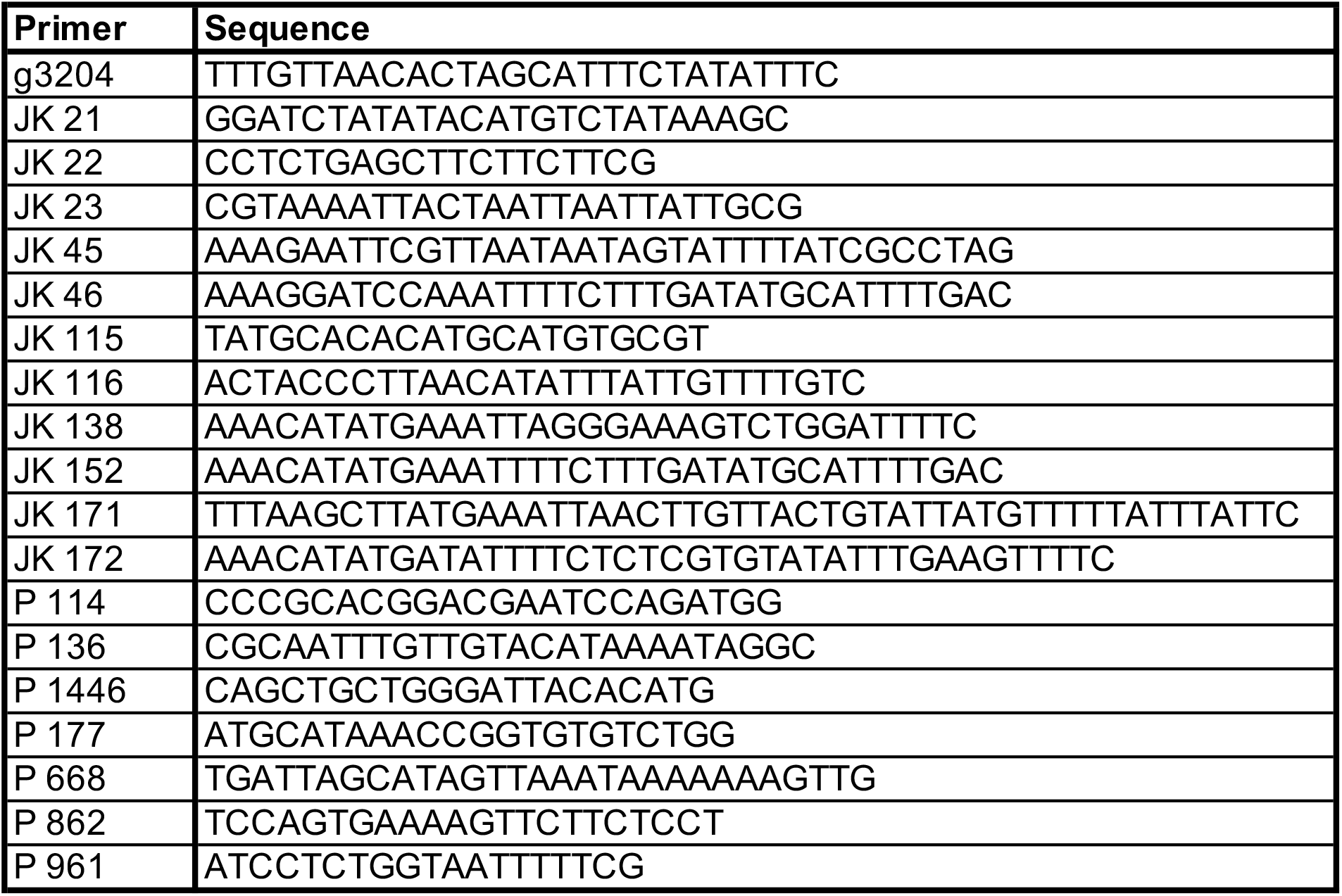
Primer sequences

